# Rapid changes in chromatin structure during dedifferentiation of primary hepatocyte*s in vitro*

**DOI:** 10.1101/2020.09.21.307363

**Authors:** Morten Seirup, Srikumar Sengupta, Scott Swanson, Brian E. McIntosh, Mike Colins, Li-Fang Chu, Zhang Cheng, David U. Gorkin, Bret Duffin, Jennifer M. Bolin, Cara Argus, Ron Stewart, James A. Thomson

## Abstract

Primary hepatocytes are widely used in the pharmaceutical industry to screen drug candidates for hepatotoxicity, but isolated hepatocytes quickly dedifferentiate and lose their mature metabolic function in culture. Attempts have been made to better recapitulate the *in vivo* liver environment in culture, but the full spectrum of signals required to maintain hepatocyte function *in vitro* remains elusive. Here we studied the dedifferentiation process in detail through RNA-sequencing of hepatocytes cultured over eight days. We identified three distinct phases of dedifferentiation. An early phase, where mature hepatocyte genes are rapidly downregulated in a matter of hours. A middle phase, where fetal genes are activated, leading to hepatocytes with a fetal phenotype. A late phase, where initially rare contaminating non-parenchymal cells overgrow the culture as the hepatocytes gradually die. Using genetically tagged hepatocytes, we demonstrate that the cells reactivating fetal marker alpha-fetoprotein arise from cells previously expressing the mature hepatocyte marker albumin, and not from albumin negative precursor cells, proving that hepatocytes undergo true dedifferentiation. To better understand the signaling events that result in the rapid down-regulation of mature hepatocyte genes, we examined changes in chromatin accessibility of hepatocytes during the first 24h of culture using ATAC-seq. We find that drastic and rapid changes in chromatin accessibility occurs immediately upon start of culture. Using binding motif analysis of the areas of open chromatin sharing similar temporal profiles, we identify several candidate transcription factors potentially involved in the dedifferentiation of primary hepatocytes in culture.

## Introduction

Hepatotoxicity can result in the termination of clinical drug trials and recall of drugs already in the market [1]. Occasionally approved drugs can cause hepatotoxicity, with more than 40,000 yearly cases of drug-induced liver injury reported in the United States alone [2]. In addition, hepatocytes can convert nontoxic compounds into compounds that are toxic to other organs, as is seen with nephrotoxicity of haloalkenes [3] and cardiotoxicity of the prodrug cyclophosphamide [4]. Animal testing is currently required before proceeding to phase I clinical trials, but the concordance to human toxicity can be poor [5, 6]. The limitations of animal models coupled with underpowered clinical studies that fail to detect rare idiosyncratic hepatotoxicity makes early discovery particularly challenging.

Primary human hepatocytes are often used in drug screening, but unfortunately they lose mature function rapidly in tissue culture [7]. Dramatic transcriptomic changes occur in as little as 30 minutes [8, 9], allowing only a short window of usefulness for toxicological screens. Mature genes such as albumin (ALB) and genes involved in hepatic xenobiotic metabolism such as CYP1A2 and CYP2E1, part of the cytochrome P450 superfamily, and gluconeogenetic enzymes including glusoce-6-phosphatase catalytic unit (G6PC) are downregulated in culture [10, 11]. Several culture techniques have been developed in an attempt to overcome this limitation [12–14], but no approach has been able to completely maintain the mature phenotype during extended culture [15]. Therefore, we hypothesis that in order to understand the full range of signals for maintaining hepatocyte maturity, it requires a detailed time course study at the molecular level.

To better understand the events that lead to the down-regulation of metabolic genes in culture, we examined changes in both gene expression by RNA-sequencing and chromatin structure by ATAC-sequencing (Assay for Transposase Accessible Chromatin with high throughput sequencing), [16] in a detailed time course study beginning immediately after the initiation of hepatocyte culture. Through our work, we identified DNA binding motifs of transcription factors potentially involved in the earliest stages of phenotype loss in hepatocyte culture.

## Results

### Primary hepatocytes undergo true dedifferentiation in culture

Primary mouse hepatocytes expressing mCherry under the control of the enhancer/promoter region of fetal hepatocyte marker alpha-fetoprotein (Afp), were isolated and cultured *in vitro* for up to 192h (RNA-seq_0h-192h dataset, supplemental table 1)[17].Initially the cells looked healthy, often observed with multiple nuclei, reflecting the *in vivo* mature hepatocyte morphology. Over time, the cell morphology appeared progressively disorganized and stringy (Figure 1A). Cultured cells were collected and mCherry expression was quantified using fluorescence-activated cell sorting (FACS). A transient phase of Afp expression was detected where mCherry expression was not seen from 0 through 36h (Figure 1B) but became detectable after 48h in culture, and the positive fraction increased to a maximum of 12.4% at 84h and stayed high until 156h after which it decreased. The emergence of Afp in culture was further confirmed in 2 separate RNA-sequencing experiments (Figure 2B and supplemental figure 4C)

**Figure 1.**
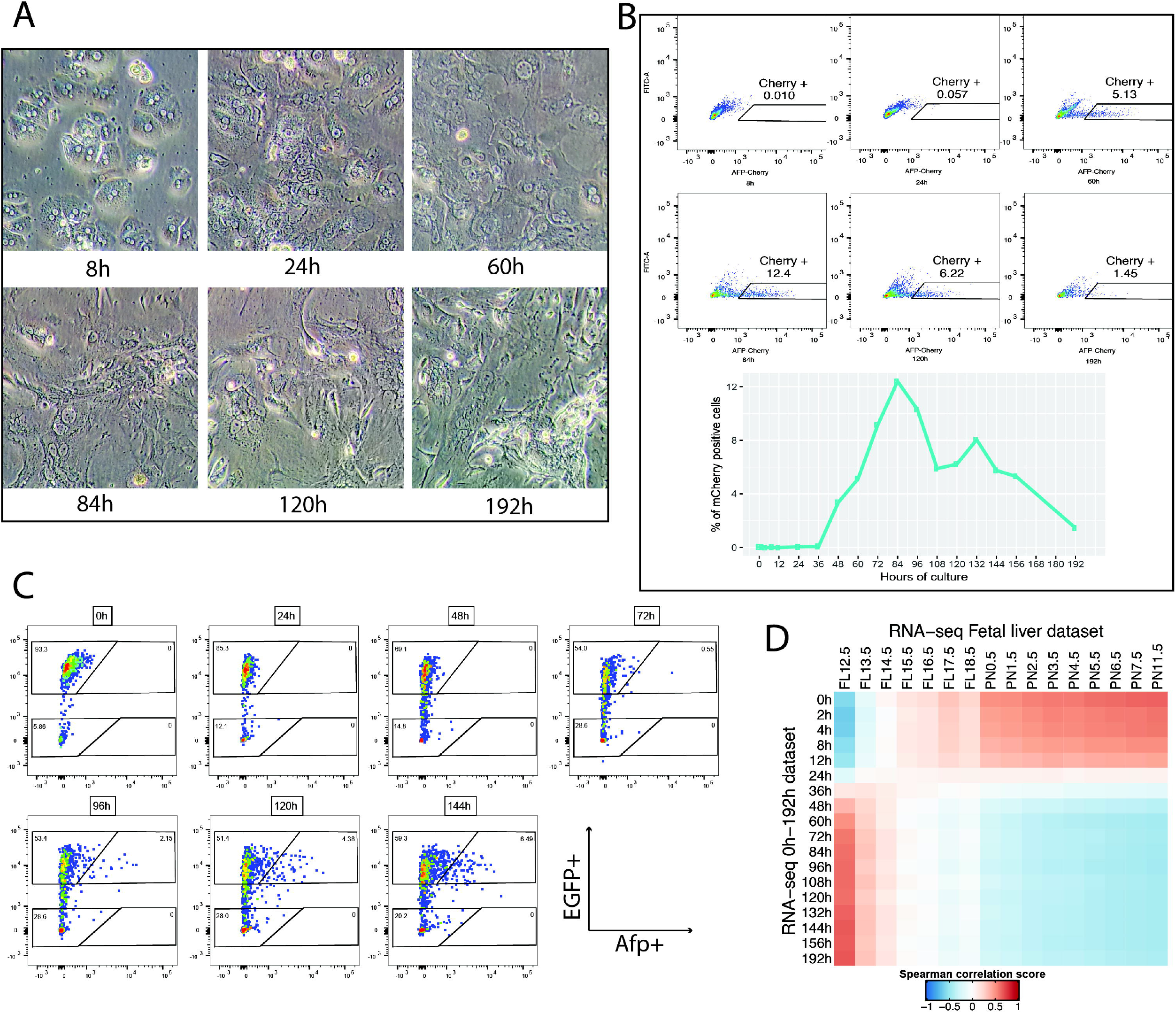
Primary hepatocytes lose their mature phenotype in culture. A) Morphology of primary hepatocytes at different times during culture. B) FACS plot identifying the emergence of an Afp+ cell population during hepatocyte culture. C) FACS plot identifying the emergence of an Afp+ cell population in culture of Alb+ hepatocytes permanently marked by EGFP+. D) Heatmap of spearman correlation between RNA-seq_0h-192h data and RNA-seq_Fetal_liver data using the top 100 upregulated and top 100 downregulated genes between 0h and 96h of the RNA-seq_0h-192h dataset.

**Figure 2.**
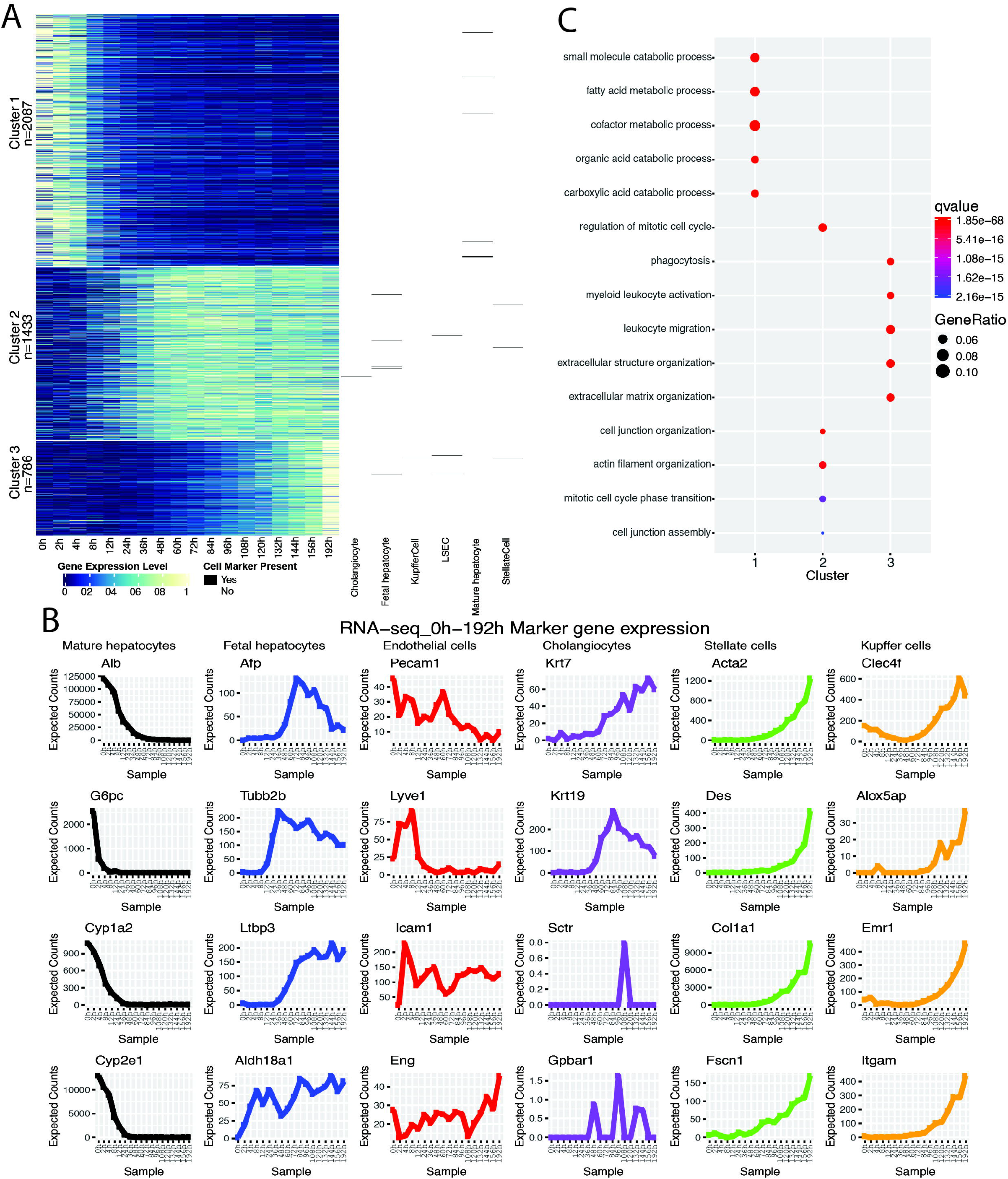
A fraction of hepatocytes in culture undergoes dedifferentiation. A) k-means clustering of the RNA-seq_0h-192h dataset with marker gene of mature hepatocytes, fetal hepatocytes, cholangiocytes, stellate cells, sinusoidal endothelial cells, and Kupffer cells placement to the right. B) GO analysis of genes from clusters in A) using the metabolic functions GO terms. C) Expression profile of marker genes for different liver cells during culture.

Because hepatocyte culture over time displays a combination of complex morphological change and proliferation, we decided to take a fate mapping approach to interrogation of whether the detection of fetal liver marker expression is due to the proliferation of rare immature cells or due to dedifferentiation of mature hepatocytes. To genetically label Albumin(Alb)-expressing hepatocytes and its progeny, a transgenic Alb promoter/enhancer-Cre driver line was crossed with a *Rosa26-loxP-stop-loxP-EGFP* Cre reporter line [18, 19]. To enable simultaneous monitor of Afp-expressing, we further crossed the Alb-Cre/R26-EGFP line with the Afp-mCherry line. Isolated hepatocytes were cultured for 144h sampling every 24h. FACS analysis (Figure 1C) showed that the population of EGFP positive cells dropped from 93% to 55% between 0–72h in culture, while the fraction of Afp-mCherry positive cells peaked at 6% after 144h. Importantly, all Afp-mCherry positive cells were also EGFP positive. Thus, our data unequivocally demonstrated that Afp-expressing hepatocytes originated exclusively from an Alb-expressing hepatocyte population.

To confirm that *in vitro* dedifferentiated hepatocytes assume a fetal phenotype, we performed RNA-sequencing of the cultured cells at the same time points used previously and calculated the spearman correlation between each time point in culture to each day in liver development from fetal day 10.5 to postnatal day 12.5 using the 100 most upregulated and 100 most downregulated genes (based on fold change) with a normalized expression above 50 expected counts (EC) at any time between 0h (most mature) and 96h (most fetal with high AFP expression) from our *in vitro* dedifferentiation time course. Among the genes included in the comparison are several markers of fetal hepatocytes such as Afp, Krt19 and Abcc1, as well as several markers of mature hepatocytes such as Cyp1a2, Cyp2e1, G6pc and Tat (Supplemental table 2). The correlation matrix (Figure 1D) shows that the most mature hepatocytes in culture (0, 2, 4, 8, and 12h) correlate poorly with the early fetal liver developmental time points but correlate progressively better with the later developmental time points. Conversely, the more dedifferentiated cells in culture (36-196h) correlate poorly to the later time points during liver differentiation but correlate progressively better to early fetal liver development time points, this is particularly evident when comparing to the earliest time point, FL12.5, demonstrating true dedifferentiation of cultured hepatocytes.

### Loss of phenotype of cultured primary hepatocytes takes place in three phases

To explore the transcriptomic dynamics of hepatocytes during culture, we clustered the transcriptomic data generated previously, using k-means clustering, after enriching the dataset for highly dynamic genes by removing the genes with less than 4 fold change throughout the culture. Different values for k were evaluated by maximizing the between sum of squares, minimizing the within sum of squares of the clusters, and penalizing for increasing number of k, resulting in a choice of k=3 (Figure 2A) (Supplemental materials and methods, Supplemental Figure 1A). Cluster 1 comprises genes with early high expression and rapid downregulated between 4–24h; cluster 2 contains genes with low early expression that increases after 8h and plateaus after 36–48h. Cluster 3 includes genes with low expression prior to 84h of culture, after which the expression increases exponentially. We mapped the location within the clusters of marker genes for mature hepatocytes, fetal hepatocytes, cholangiocytes, stellate cells, liver sinusoidal endothelial cells, and Kupffer cells on the heatmap. All mature hepatocyte markers were found in cluster 1, demonstrating that hepatocytes lose their phenotype after a short time in culture. The majority of the fetal hepatocyte markers were found in cluster 2, while most of the marker genes for stellate cells and Kupffer cells were found in cluster 3 and were highly expressed in the later time points. Most cholangiocyte and liver sinusoidal endothelial cell marker expression were low, excluding them from the heatmap, further no trend was found in those genes that were included.

Mature hepatocyte markers, such as G6pc (glucose-6-phosphatase, catalytic) and albumin, are quickly downregulated (Figure 2B first column), and expression becomes undetectable after 8–12h and 24–36h of culture respectively. Most of the phase 1 and 2 genes (e.g. Cyp1a2 and Cyp2e1) fall in between these time points. Markers of fetal hepatocytes (Figure 2B second column) were found to appear as early as 2–4h (e.g. Aldh18a1), while Tubb2b came up after 12–24h of culture. Common fetal markers Afp and Ltbp3 were expressed after 36–48h. Interestingly, we found Alb and Afp expression levels to display opposing profiles, with Afp expression increasing shortly after Alb expression declines. This pattern mimics, in an inverted fashion, the downregulation of Afp followed by upregulation of Alb seen in neonatal livers, suggesting a common opposing regulatory mechanism [20]. Nonparenchymal cell markers had no consensus (Figure 2B column 3). However, Pecam1, an endothelial cell marker decreased during the culture. Cholangiocyte markers Krt7 and Krt19 (Figure 2B column 4) showed a pattern similar to that of fetal hepatocyte markers. Possibly recapitulating their expression in hepatoblasts, a common precursor for both hepatocytes and cholangiocytes. No other markers of cholangiocytes were expressed, suggesting the observed keratin expression to be due to dedifferentiation of hepatocytes. Finally, markes of both stellate cells (Figure 2B column 5) and Kupffer cells (Figure 2B column 6) increased exponentially toward the end of the time course.

Examining hepatocytes permanently marked by enhanced yellow fluorescent protein (EYFP) using FACS (RNA-seq_0h-169h_EYFP dataset, detailed description in Supplemental table 1) (Supplemental Figure 4C), we found a similar expression pattern for mature and fetal hepatocyte markers, although the response was delayed likely due to slight changes in viability and thus final cell density. The expression of stellate and Kupffer cell markers on the other hand is greatly decreased or not detected.

We analyzed the genes found in the three clusters of Figure 2A using gene ontology (GO) analysis. The top five enriched terms for cluster 1 consist of metabolic functions consistent with mature hepatocyte function (Figure 2C). Cluster 2 is enriched in terms related to cell division and cell junction organization, consistent with the hepatocytes attempting to proliferate and regrow the liver as hepatocyte isolation possibly mimics liver injury. Finally, cluster 3 is enriched in terms related to leukocytes and extracellular matrix, consistent with the dominance of extracellular matrix secreting activated stellate cells and Kupffer cells, the residential macrophages of the liver.

Taken together, the data suggests that changes in cultured primary hepatocytes happens in three phases. The first phase is a rapid general down regulation of genes associated with mature hepatic phenotype. This is followed by a second phase where the mature hepatocytes dedifferentiate and initiates expression of fetal hepatocyte marker genes. Finally, in phase 3, the data suggests Kupffer cells and activated stellate cells, a small fraction of the initial population, divides faster than hepatocytes and outnumber them toward the end of the time course.

### Primary hepatocytes in culture undergo rapid epigenetic changes

We hypothesized that the key events that lead to the loss of mature phenotype/dedifferentiation of primary hepatocytes in culture happens within the first 24h, at which point fetal hepatocyte markers are already being expressed (Figures 2A and 2B). We therefore decided to investigate the initial 24h of primary hepatocyte culture in greater detail (Figure 3A) and obtain both global chromatin accessibility status (ATAC-seq) and transcriptomic (RNA-seq) data. To avoid examining an increasingly heterogenous cell population, we sort out EGFP positive cells from the Alb-Cre/R26-EGFP transgenic mouse model, allowing us to exclusively analyze cells belonging to the original hepatocyte population.

**Figure 3.**
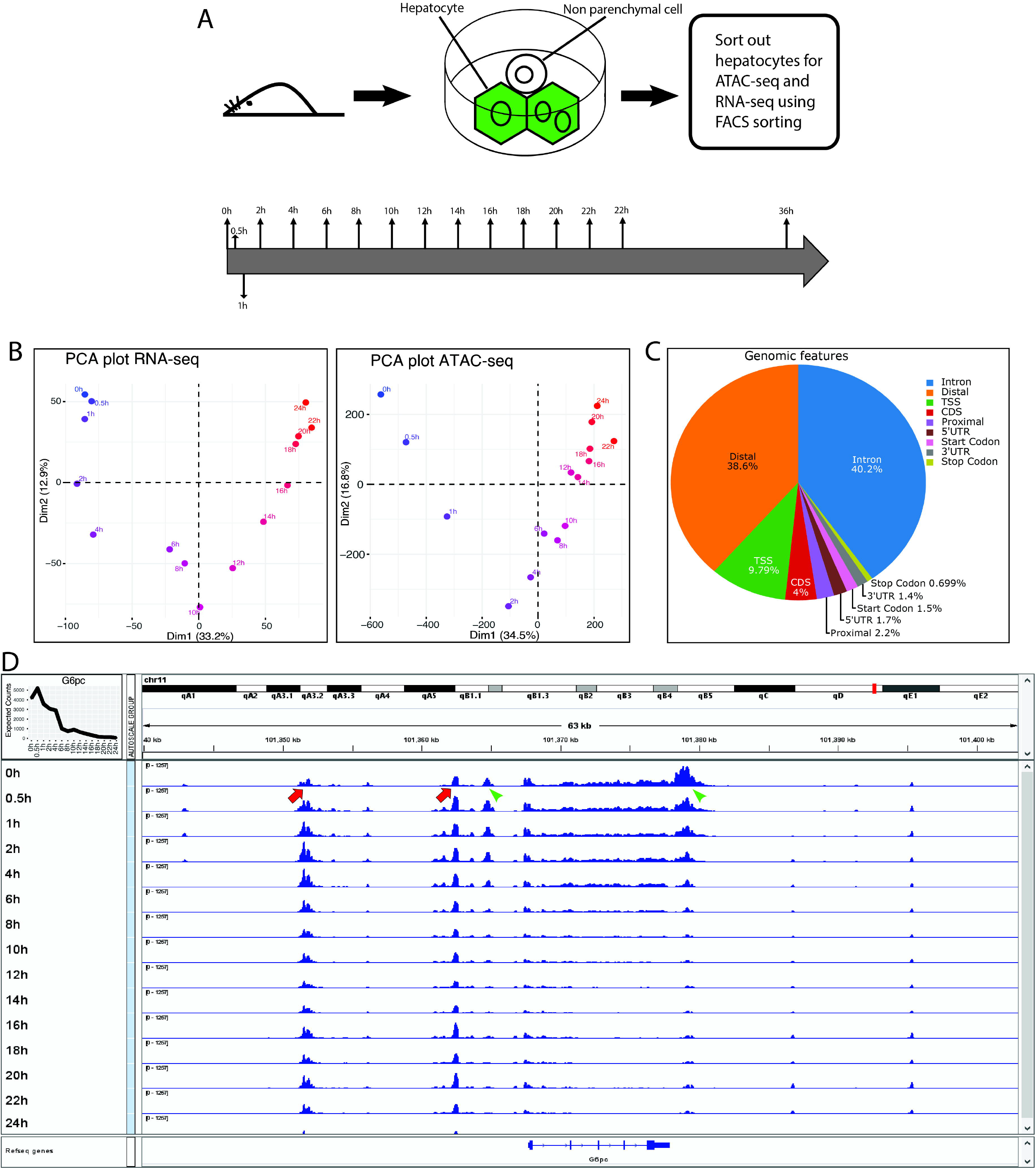
Overview of chromatin accessibility during 36h of culture. A) Overview of experiment. B) Plot showing PCA dimension 1 and 2 of gene expression (left) and open chromatin (right) across 24h of culture. The color gradient ranges from 0h of culture (blue) to 24h of culture (red). C) Fraction of areas of open chromatin found in genomic features. D) Genome browser visualization of the area around the G6pc gene (transcribed from left to right). Blue peaks represent open chromatin. 0h of culture is found at the top of the figure with time progressing to 24h at the bottom. The insert on the top left displays the RNA expression level of G6pc. Red arrows highlight static peaks and green arrowheads highlight dynamic peaks.

Principal component analysis (PCA) analysis of the two datasets (Figure 3B) shows that for both RNA-seq (RNA-seq_0h-24h_EGFP, supplemental table 1) and ATAC-seq (ATAC-seq_0h-24h_EGFP, supplemental table 1) PCA 1 and 2 give a clear separation of the time points in a sequential temporal pattern. In the RNA-seq data the first 3 time points (0–1h) cluster together, suggesting only minor changes, whereas the 1–6h time points display the highest dynamic changes and we find a gradual change until 18h after which, the time points cluster together again. The ATAC-seq data displays major dynamic changes between 0-2h. The changes are found to be gradual from 2-12h, after which the timepoints cluster together suggesting less dynamic change between these timepoints.

Examining the genomic features of the area of open chromatin (AOC) (Figure 3C), we find that 38.6% of all the AOCs are found in distal areas (>1kb upstream from the nearest transcription start site or TSS) of the genome, 2.2% are found in the proximal (<1kb upstream from the nearest TSS) area, 9.79% are found to span the TSS and 49.5% of the AOCs are found inside the gene. Interestingly, at 40.2% most of the peaks found inside the genes were found in intronic regions.

Using a genome browser, we examined the chromatin landscape around G6pc (Figure 3D), one of the fastest downregulated mature hepatocyte markers. It is apparent that there are several AOCs around this locus. Not all display a dynamic profile; two examples of static AOCs (highlighted with red arrows) display little relation to the gene expression profile, whereas two examples of dynamic AOCs (highlighted with green arrows) display a profile similar to the expression of the gene. This suggests that not all AOCs regulate the expression of the nearest genes. A key issue with ATAC-seq data sets is to identify the signal among the largest number of AOCs identified. By focusing exclusively on AOCs with a dynamic temporal profile, we enrich for AOCs presumably involved in regulating the expression of genes with a similar dynamic expression profile, thereby increasing the signal and lowering the noise in the dataset.

### Changes in chromatin accessibility happens in six distinct patterns

We identified a total of 192,621 AOCs, with a mean of 95 cutting events (CE) per area. A CE denotes an end of a mapped fragment. To reduce noise, we filtered out AOCs with less than 95 CE at any time point to remove the 50% with the lowest number of CE. We selected dynamic AOCs with a log2 fold change of 2.3 (~5-fold) or higher in CE across the time course. This left a set of 11,184 AOCs with a majority located within a 100kb area of the TSS (Supplemental Figure 2A). The distribution of the AOCs were skewed toward the upstream side of the TSS and a distinct binding pattern 1kb upstream and downstream of the TSS were identified. We also found increased accessibility around randomly selected gene locations, but a clear difference was seen when we subtracted random signal from the true signal (Supplemental Figure 2C, Supplemental materials and methods).

Similar to the approach taken with genes in the RNA-seq data, we grouped AOCs using k-means clustering on normalized CEs across the time course. By optimizing the sum of squares ratio (Supplemental materials and methods), we identified six k-means clusters to best represent the dataset (Supplemental Figure 1). Examining the heatmap of these seven clusters (Figure 4A left) we find that the AOCs in cluster 1 are highly accessible initially but disappear quickly during the first hour of culture. Cluster 2 consists of inaccessible areas at timepoint 0 hour that rapidly become accessible at the 0.5h timepoint and is accessible until the 1h after which they disappear. Cluster 3 is seen to be mostly inaccessible at 1 hour and gains quickly maximal accessibility after 2h before gradually becoming less available toward the latter time points. The chromatin in cluster 4 become accessible after 2-4h of culture and maintains accessibility for the rest of the time in culture. Cluster 5 becomes accessible starting at 2-4h and increasing until 14-16h of culture after which the accessibility is maintained. Finally, cluster 6 is found to become accessible around 16-18h of culture and increases for the duration of the time course. The high temporal sampling rate allowed the sensitivity needed to detect both the highly dynamic AOCs, such as seen in clusters 2 and 3 and the speed of loss of chromatin accessibility in cluster 1, which otherwise might have been missed. The changes to the chromatin accessibility seen in clusters 1 and 2 occur within the first half hour after culture, and although protein expression can be detected as early as early as 10 to 20 min after stimuli [18], very few proteins in mammals have a half-life less than an hour [19], making it possible that these areas of accessible chromatin are regulated non-transcriptionally, such as by phosphorylation of TFs, and would be undetectable by RNA-sequencing alone.

**Figure 4.**
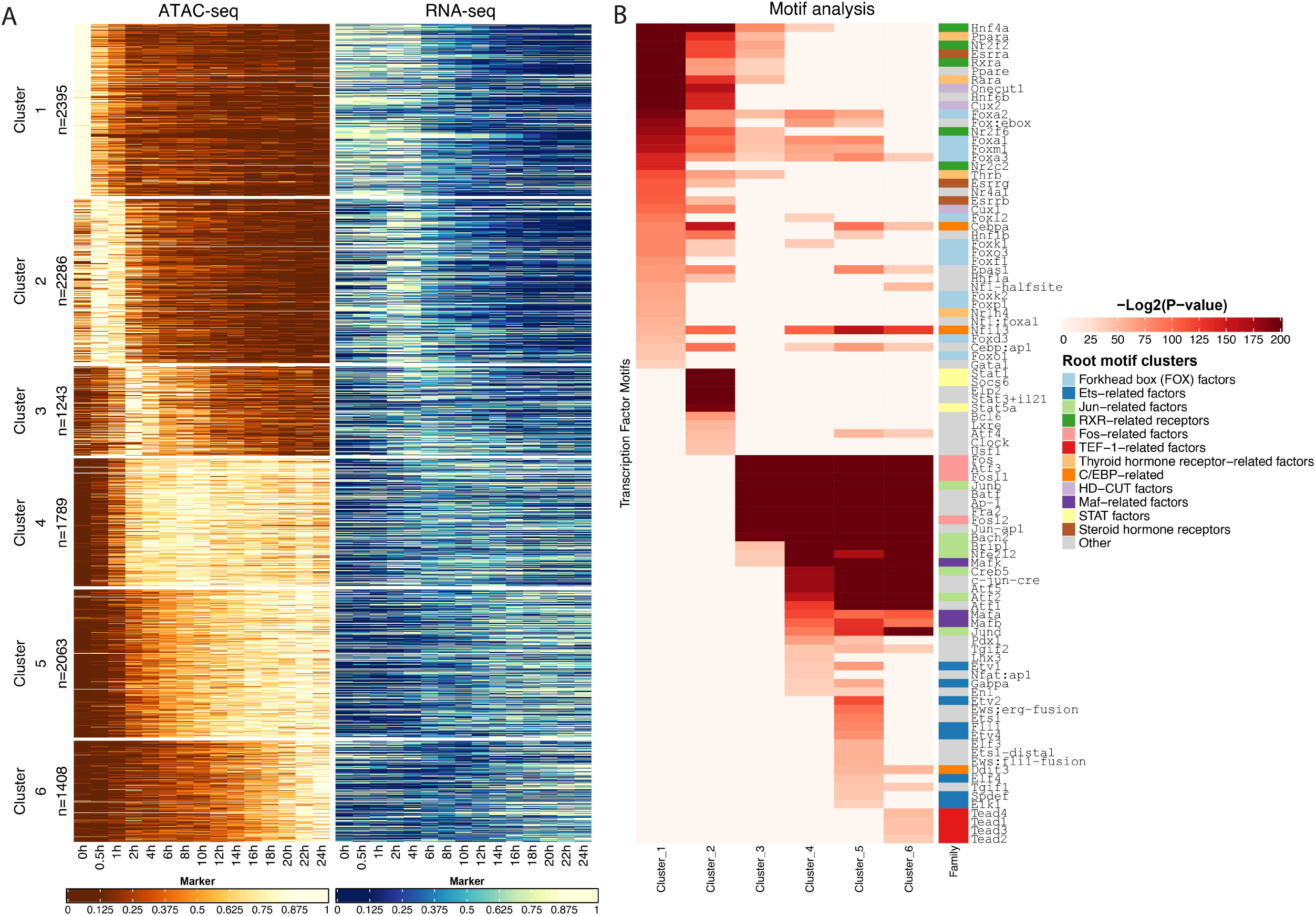
Open chromatin of primary hepatocytes changes rapidly during cell culture. A) K-means clustering of open chromatin across the time course (left) with complementary gene expression profile of the gene with the nearest TSS to the area of open chromatin in the left heatmap (right). The gene expression heatmap (right) is ordered based on the nearest AOC – the nearest gene and AOC is in the same row. B) Enriched TF motifs in open chromatin of each clusters from A). The colored column depicts TFs belonging to the same family. The enriched TF is displayed on the right-hand side of the heatmap.

### RNA expression levels are delayed and correlates with open chromatin

We created a heatmap of gene expression by assigning the gene with the nearest TSS to each AOC and created a heatmap using the clustering order established for ATAC-seq data, so the AOC and the gene with the nearest TSS is placed in the same row of both heatmaps (Figure 4A left and right). We observed that when a region near a TSS is accessible, the gene is typically expressed and vice versa. This is most evident for clusters 1 and 2. This association supports the idea that accessibility of chromatin and RNA expression levels of genes around them are linked. Interestingly, we find that the change in RNA expression is delayed compared to the changes seen in chromatin accessibility. This is seen clearly in clusters 2 and especially cluster 3, where the upregulation of gene expression is delayed for about 2h following the opening of the chromatin[21].

### Identification of TFs involved in dedifferentiation of hepatocytes

To identify putative causal agents of the changes seen in the AOC profiles, we decided to examine the involvement of TFs. We assumed that AOCs with a similar dynamic profile were likely regulated by the same group of TFs. We used the DNA sequences of the AOC in the seven clusters from Figure 4A (left) to test for enrichment of known TF binding motifs. The result of the motif enrichment analysis is shown in Figure 4B, which includes all motifs with an adjusted p-value of less than 1×10^−10^ for each cluster. The AOCs from cluster 1 which rapidly closes and becomes inaccessible in culture and is enriched in TFs belonging to the RXR-related receptors family, such as Hnf4a, Nr2f2, Rxra, and the Fox-factor family, such as Foxo3, Foxa1, and Foxa2. The RXR-related receptor motifs are also found enriched in clusters 2 and 3 but less significantly. Further, the Fox factor family was found to be enriched in AOCs from other clusters but less significantly than in cluster 1.

Cluster 2 was enriched for the Stat-factor TF family, with Stat1, Stat3, and Stat5 being specifically enriched. Clusters 3 through cluster 6 were found to be increasingly enriched of Fos-, Jun- and Maf-related factor families, such as Fosl1, Junb, and Mafk, which was found to have adjusted p-values in the orders of 10^−1000^. Further in cluster cluster 5 motifs from members of the Ets-related factors family (Etv2, Etv1, Elk1 etc.) are found to be enriched, and in cluster 6 we find enriched motifs belonging to theTef1-related factors (Tead1, Tead2, Tead3 and Tead4).

## Discussion

The use of primary hepatocytes for toxicological testing is limited due to their rapid loss of mature phenotype in culture, a process described as dedifferentiation in the literature [20]. Here we show that while the entire population of hepatocytes in culture loses the mature phenotype, a small subset of the cells undergo true dedifferentiation and activate expression of the fetal hepatocyte marker Afp after 48h, paralleling the *in vivo* response to partial hepatectomy [21]. We found that 6-10% of the hepatocytes in culture activate Afp expression, and that the population expressing Afp derives exclusively from an Alb expressing hepatocyte pool. Further, after 48h of culture, the transcriptomes of hepatocytes showed a high correlation to the early stages of liver development, whereas prior to 24h they correlated better to postnatal livers. In short, longer culture periods took hepatocytes further back in the development process.

Dedifferentiation was found to happen in three phases, an early phase marked by downregulation of genes specific to mature hepatocytes, resulting in loss of mature phenotype; a middle phase with an initial rapid change in gene expression and chromatin accessibility as the hepatocytes respond to their new environment, followed by expression of fetal marker genes; and a late phase, where other cell types, such as activated stellate cells and Kupffer cells, dominate the culture.

Genome-wide chromatin accessibility and complementary gene expression data over temporally dense sampling times of hepatocytes cultured for 24h allowed us to investigate early causal factors of hepatocyte dedifferentiation. Major changes in AOCs were found after as little as 30 min in culture and assigning each AOC to the gene with the nearest TSS revealed a clear relationship between the dynamics of AOCs and gene expression. In some cases, we detected a lag in observed changes to transcription versus corresponding changes in AOCs. This was especially evident in AOCs linked to gene activation. It is known that changes in chromatin accessibility precede changes in gene regulation [22], and the high temporal resolution of this dataset allowed us to identify a ~2h time shift between changes in AOCs and gene expression changes within some clusters.

Interestingly, by clustering the AOCs, we found that groups regulated in a similar fashion temporally, were also enriched for similar TF binding motifs. We found a strong enrichment of motifs from the Fos and Jun families in AOCs after 2h of culture (clusters 3–6). This family is important for proliferation, differentiation, survival, and apoptosis [23]. AOCs at later time points (cluster 6) are enriched for the Tef-1-related factor family, such as the Tead TFs. Tead TFs are known to play a role in determining the size of the liver [24] and maintaining a mature phenotype in hepatocytes [25].

Regulatory changes in the first hour of culture are of special interest, as these initial events should help identify the signals that sustain mature hepatocyte function in vivo but are missing in vitro. Motifs for TFs belonging to the Rxr-related family, such as Hnf4a and Rxra, are enriched in AOCs that quickly become inaccessible (Clusters 1 and 2). The factors in this family are known to regulate mature hepatocyte genes [26–28]. Other TFs with known functions in the mature liver [29, 30] were also identified in these clusters, such as Hnf1a, Hnf1b, and several Fox TF family members. Further, TFs belonging to the Stat family were found to be enriched in cluster 2, where the chromatin is transiently accessible from 0.5h-1h. The Stat TFs, part of the Jak-Stat pathway, play an important role in inflammation, proliferation, differentiation, and apoptosis [32, 33], and have previously been linked to maintenance of mature phenotype in hepatocytes [34, 35].

The findings here indicate that a fraction of hepatocytes in culture undergo true dedifferentiation accompanied by expression of fetal marker genes and increased correlation to early liver development. Most cells however lose expression of mature genes without activation of fetal genes.

Using genome-wide chromatin accessibility data combined with complementary gene expression profiles at a high temporal resolution allowed us to examine the changes in chromatin accessibility and identify several key TF families involved in the early dedifferentiation of cultured hepatocytes. Further this dataset creates a valuable resource for researchers exploring the dynamics of epigenetic and gene expression changes of hepatocytes *ex vivo, and thereby* aiding the development of protocols to maintain maturity of primary hepatocytes in culture.

## Materials and Methods

### Animals and handling

All animals were kept under standard husbandry conditions. Details on mice used are given in Supplemental Table 1. Animal experiments and procedures were approved by the University of Wisconsin Medical School’s Animal Care and Use Committee and conducted in accordance with the Animal Welfare Act and Health Research Extension Act.

### Hepatocyte isolation

Hepatocytes were isolated using a two-step perfusion protocol [22] Mice were euthanized by isoflurane followed by cervical dislocation. Liver Perfusion Medium and Liver Digest Medium (Gibco) were perfused for 10 minutes at 7 ml/minute flowrate. Hepatocytes were purified by centrifugation at 50 x G, 4 times for 3 minutes, discarding the supernatant.

### Cell culture

Hepatocytes were cultured under standard conditions (37°C., 20% O_2_, and 5% CO_2_), with 10^6^ viable cells/well in a 6-well plate precoated with matrigel in Williams E medium (Thermo Fisher # 12551-032) + 1x ITS (Thermo Fisher # 41400045) + 1x Glutamax (Thermo Fisher # 35050061) + 10% FBS (Thermo Fisher # 16000044). After 2 hours in culture, the FBS was removed. Media was changed every 24hours.

### Datasets

Datasets used are detailed in Supplemental Table 1.

### RNA-sequencing data analysis

All RNA-Seq sequences were filtered and trimmed to remove low quality reads, adapters, and other sequencing artifacts. Reads were mapped to the NCBI mm10 (GRCm38) mouse reference with Bowtie (v1.2.2). Expected counts for gene expression was calculated by RSEM (RNA-seq by expectation maximization) (v1.2.3)[23].The raw expected counts estimates were then renormalized (after removing mitochondrial genes) using the median-by-ratio algorithm bundled with the R package EBSeq (v.1.26.0) differential expression package. Normalized expression data was filtered to include only genes with >= 50 EC in at least one sample, and >=4-fold change between its minimum and maximum unless otherwise stated.

### Cell sorting

Hepatocytes were singularized using trypsin and sorted using an ARIA III cell sorter (BD Biosciences) based on the marker genes expressed in the cells.

### ATAC-sequencing data analysis

All samples sequenced on the Illumina HiSeq-3000 were demultiplexed with the illumina bcl2fastq 2.17 converter. FASTQ output from the sequencer was processed all the way through peak-calling using the epigen-UCSD ATAC-seq pipeline, which is forked from a pipeline created by A. Kundaje et al. (https://github.com/kundajelab/atac_dnase_pipelines). The pipeline was configured via the true-reps switch to run on replicate pairs corresponding to time points, with peaks called on each pair. The output peaks were post-processed, merging all overlapping peaks called across all samples, then trimming each merged peak to +/− 250bp from its summit, calculated by summing all aligned tags from all samples. (Note that duplicate tags were removed from each sample, but not across samples; these are the tag alignments found in the various atac_bds output files of the form XXX.trim.PE2SE.nodup.tn5.tagAlign.gz.)

The merged and truncated peaks were then rescored for each replicate-pair, summing over 25 million randomly selected aligned tags from each replicate. The resulting table of counts per peak for each replicate pair was then renormalized using the quantile normalization algorithm in the R preprocessCore package (v.1.48.0). These renormalized tables of peak-counts across time points were then used for subsequent investigation.

### Heatmaps

All heatmaps were created using the ComplexHeatmap (v.2.2.0) package in R. For all heatmaps, each row was normalized to have a range of 0 to 1 before the data was k-means clustered using kmeans function (R package stats v.3.6.2) and drawn.

### Annotate open chromatin to nearest gene

Open areas of chromatin were annotated to genes using the annotate peak feature in HOMER (default settings, mm10 as genome, HOMER v.4.11.1)

### Gene ontology analysis

Gene ontology analysis was performed using the R package clusterProfiler, with the unfiltered gene list as background, p-value cutoff of 0.01, and a q-value cutoff of 0.05 (v.3.14.1). Only the GO terms in “Metabolic Functions” were used.

### Correlation analysis

Top 100 most up-downregulated genes between 0–96h of the RNA-seq_0h-192h dataset were selected, and a correlation table between all possible pairs across the RNA-seq datasets was calculated using R.

### Principal component analysis

PCA plots were created using the factoextra (v.1.0.6) package in R. Normalized EC from the RNA-seq_0h-24h_EGFP dataset and normalized cutting events from the ATAC-seq_0h-24h_EGFP dataset were used as input. Genes/areas with 0 variance across the time course were removed.

### Genomic feature graph

Genome features of accessible areas of chromatin were extracted from the Ensmbl annotation (Mus_musculus.GRCm38.98) with all areas within 1,000bp upstream of a transcription start site (TSS) classified as a proximal area. The fraction open chromatin on each genome feature was calculated and plotted using the R package plotly (v.4.9.2).

### Genome browser image

Bam files of the ATAC-seq_0h-24h_EGFP dataset and RNA-seq_0h-24h_EGFP dataset were converted into a bigwig file using first genomecov (bedtools v.2.29.2) to generate a bedgraph file and then bedGraphToBigWig (UCSC) and visualized using the IGV genome browser.

### Transcription factor (TF) motif analysis

Enriched TF motifs were identified using HOMER (default settings, v.4.11.1). Motifs with an adjusted p-value of 10^−10^ or lower in any peak were included. The selected TF motifs were then grouped into families using TFclass. The motifs were sorted according to adjusted p-value and plotted using complexHeatmap (v.2.2.0).

