## Supplemental material for "Rapid changes in chromatin structure during dedifferentiation of primary hepatocyte*s in vitro*"

**Supplemental Materials**

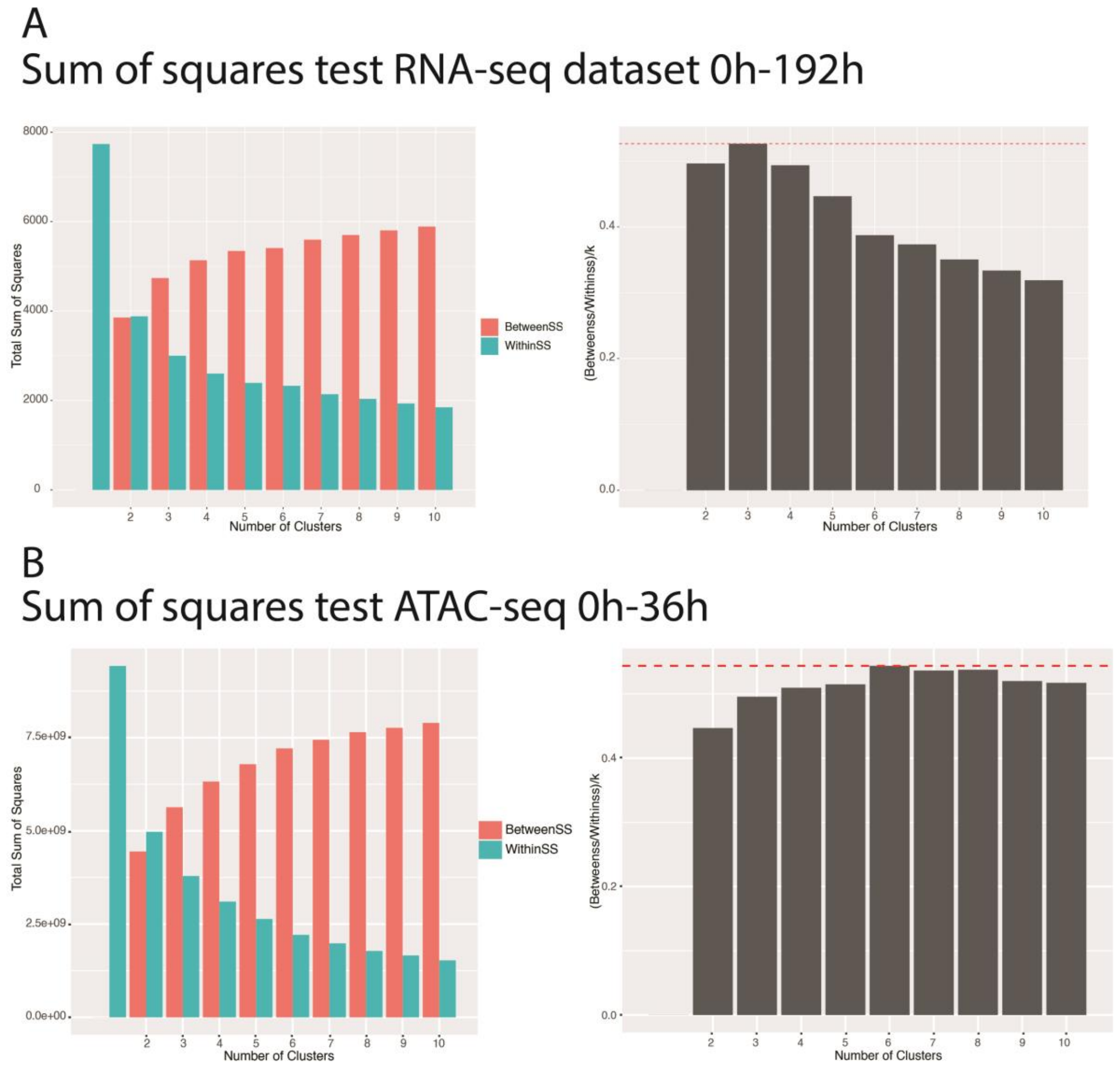


**Sup. Fig. 1. Identifying idea number of clusters for k-means clustering.**

A left) Graph is showing the values for between sum of squares (red) of clusters in k-means clustering and within sum of squares (blue) of EC/CE of genes/areas of open chromatin within clusters over 1 to 10 clusters (x-axis) for the RNA-seq_0h-192h dataset used in Figure 1. A right). The ideal number of k was identified by maximizing the between sum of squares and minimizing the within sum of squares and penalizing for increasing number of k. B) The same as A) but for the RNA-seq_0h-169h_EYFP dataset used in Figure 4).

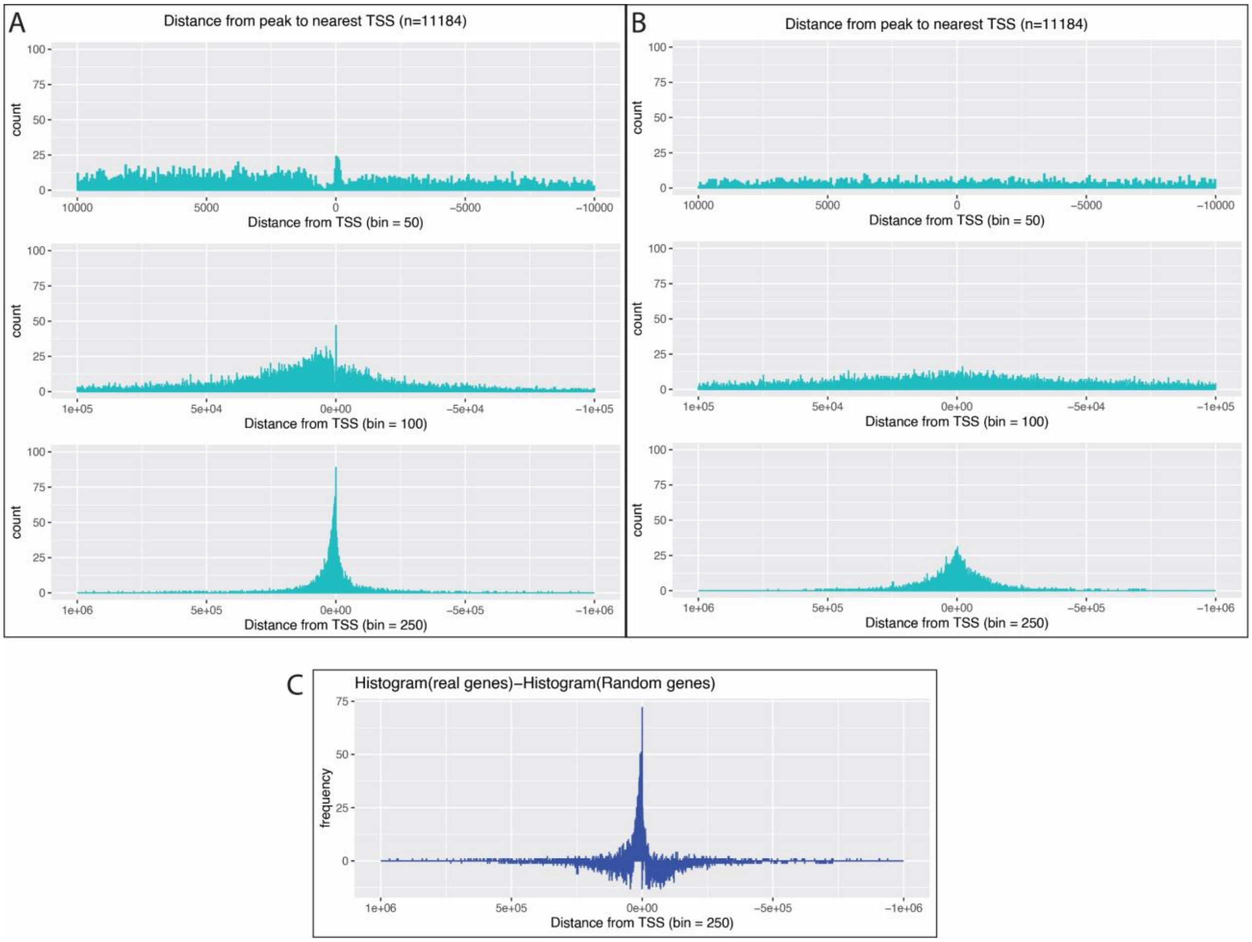


**Supplemental Figure 2. Distance from peak to TSS.**

A) Histograms showing the distance from AOC to the nearest TSS. The top graph depicts a 20kb window around the TSS, the middle graph depicts a 200kb window around the TSS, and the bottom graph depicts a 2,000kb window around the TSS. B) same as A) but showing the distance to an equal number of randomly positions on the genome. C) Shows the bottom histogram of A) with the bottom histogram of B) subtracted to highlight the differences and show that there is still a signal above background.

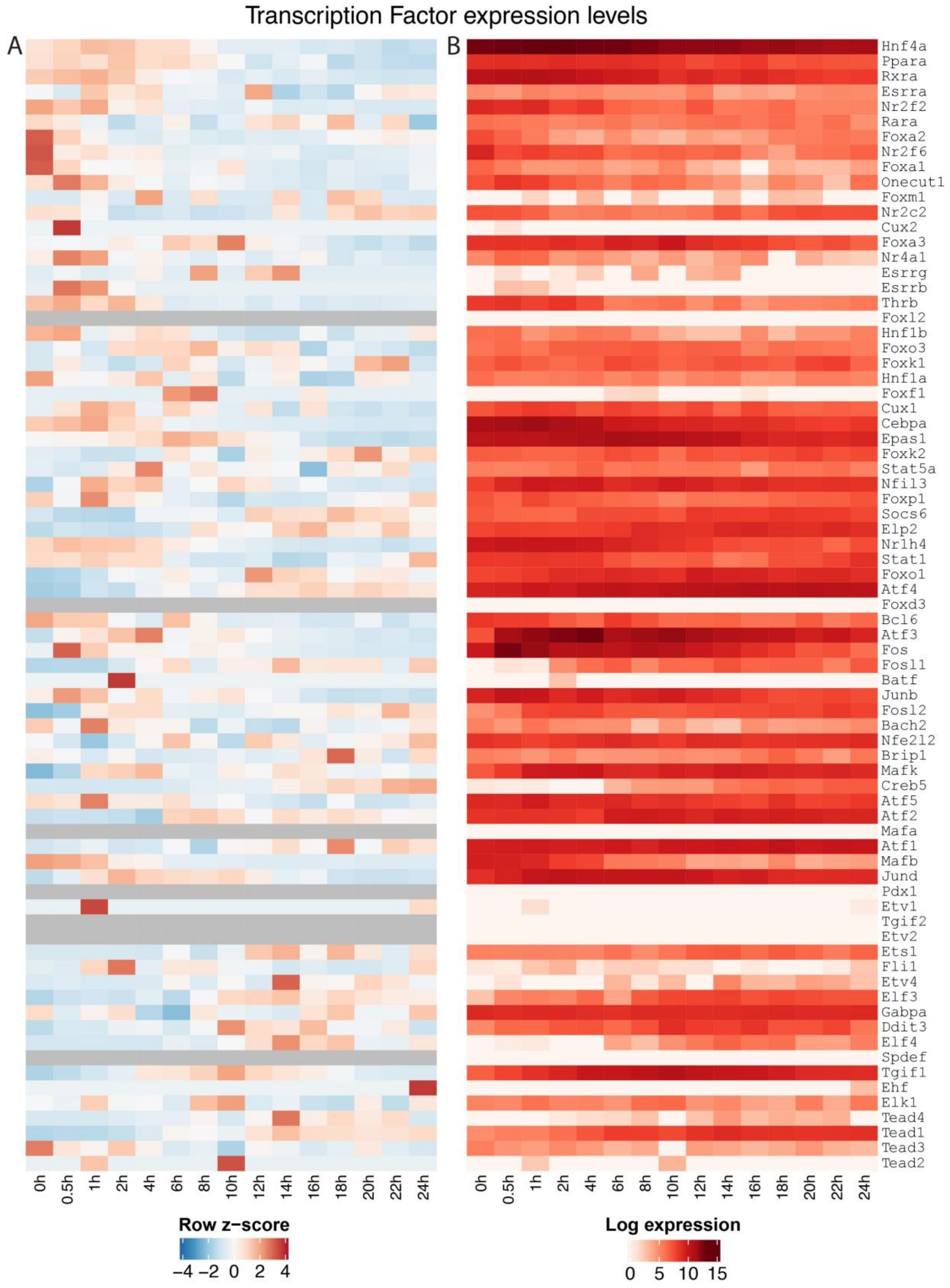


**Supplemental Figure 3. Expression profile of TFs.**

Expression of TFs found to be enriched in Figure 3. A) Row z-score of expression across the time course. B) Same as A) but using Log2 expression level.

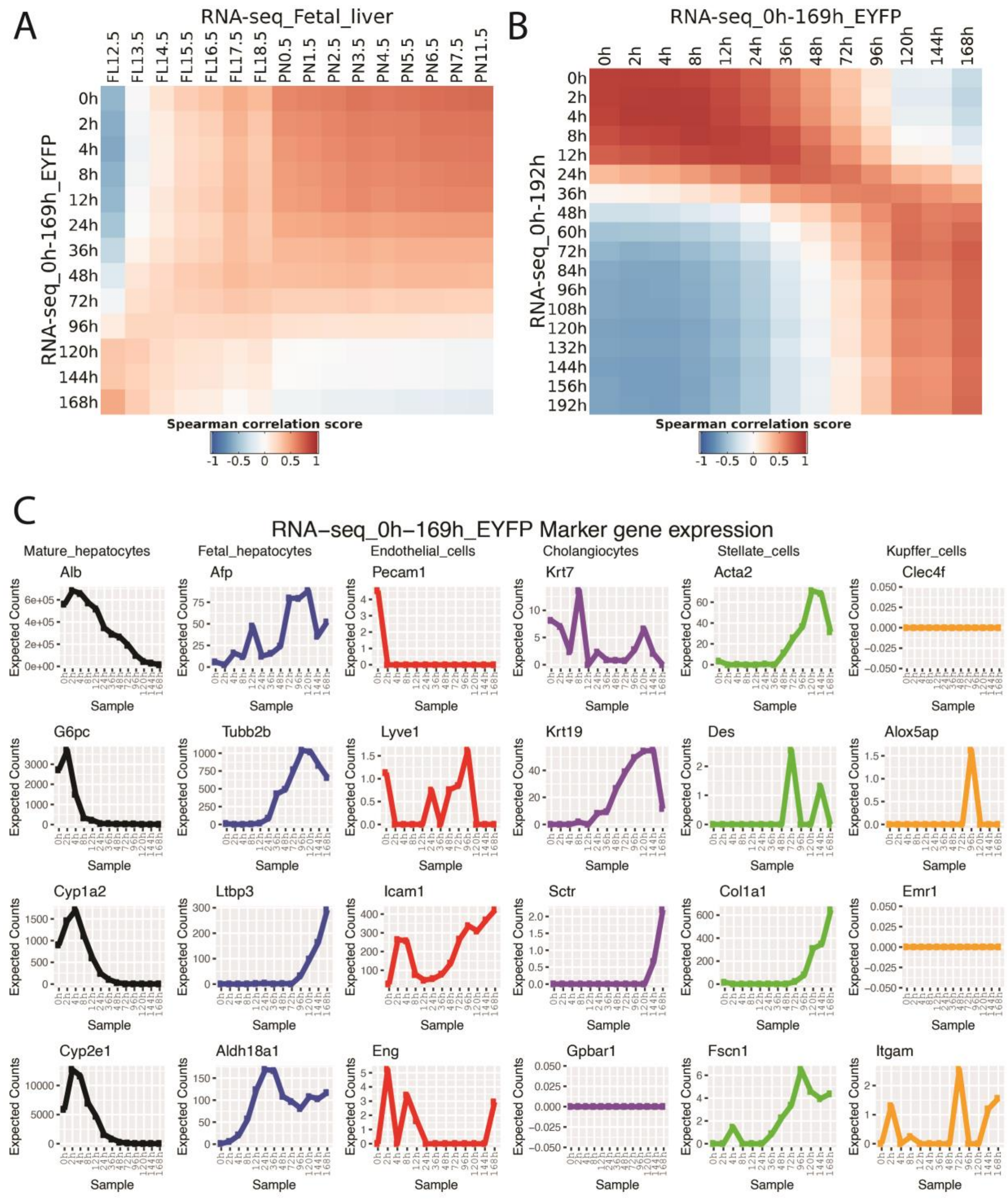


**Supplemental Figure 4. Correlation of datasets and markergene expression in EYFP sorte hepatocytes**

A) Spearman correlation comparing cultured hepatocytes marked with EYFP and sorted before sequencing with the liver developmental time course on the basis of the same genes used for Figure 2D. B) Same as A) but comparing the two primary hepatocyte culture time courses. C) Marker gene expression in EYFP-sorted dataset.

**Supplemental table 1**

| Name | Mouse strain | Type | Comments |
|---|---|---|---|
| RNA-seq_0h-192h dataset | Afp::mCherry (Jackson Laboratories stock # 014177) | RNA-seq | Mouse strain contains transgenic mCherry controlled by the Afp promotor/enhancer elements.<br>mRNA was harvested at 0h, 2h, 4h, 8h, 12h, 24h, 36h, 48h, 60h 72h, 84h, 96h, 120h, 144h, 156h, and 192h |
| RNA-seq_0h-169h_EYFP dataset | Rosa26-EYFP/Alb-CRE (Jackson Laboratories stock # 003574 x Jackson Laboratories stock # 006148) | RNA-seq | Mouse strain has loxP-stop-loxP-EYFP inserted into the Rosa26 location and a transgenic CRE recombinase controlled by Alb promotor/enhancer element permanently expressing EYFP in Alb+ cells.<br>EYFP+ cells were sorted out and mRNA was harvested at 0h, 2h, 4h, 8h, 12h, 24h, 36h, 48h, 72h, 96h, 120h, 144h, and 168h |
| RNA-seq_0h-24h_EGFP dataset | Rosa26-EGFP/Alb-CRE | RNA-seq | Mouse strain has loxP-stop-loxP-EGFP inserted into the Rosa26 location and a transgenic CRE recombinase controlled by Alb promotor/enhancer element permanently expressing EGFP in Alb+ cells.<br>EGFP+ cells were sorted out and mRNA was harvested at 0h, 0.5h, 1h, 2h, 4h, 6h, 8h, 10h, 12h, 14h, 16h, 18h, 20h, 22h, and 24h |
| ATAC-seq_0h-24h_EGFP dataset | Rosa26-EGFP/Alb-CRE (Jackson Laboratories stock # 003574 x Jackson Laboratories stock # 32037-JAX) | ATAC-seq | Mouse strain has loxP-stop-loxP-EGFP inserted into the Rosa26 location and a transgenic CRE recombinase controlled by Alb promotor/enhancer element permanently expressing EGFP in Alb+ cells.<br>EGFP+ cells were sorted out and the ATAC-seq protocol was performed at 0h, 0.5h, 1h, 2h, 4h, 6h, 8h, 10h, 12h, 14h, 16h, 18h, 20h, 22h, and 24h |
| RNA-seq_Fetal_liver dataset | C57BL/6 (Jackson Laboratories stock # 000664) | RNA-seq | Pieces of whole livers from mice at different ages were harvested and mRNA was purified. Using timed matings, livers of fetal mice were harvested at embryonic day 12.5, 13.5, 14.5, 15.5, 16.5 ,17.5 and 18.5 and at day 0.5, 1.5, 2.5, 3.5, 4.5, 5.5, 6.5, 7.5, and 11.5. |
| EGFP/mCherry lineage tracking | Afp::mCherry/EGFP/ Alb-CRE mouse (Jackson Laboratories stock # 003574, Jackson Laboratories stock # 014177 and Jackson Laboratories stock # 32037-JAX) | FACS sorting | Lineage tracing of hepatocytes using Alb-CRE and loxP-stop-loxP-EGFP. Cells are further transgenic for mCherry controlled by Afp promoter/enhancer. Cells were singularized using trypsin before being analyzed for EGFP and mCherry expression. |

| Top_1_to_50_upregulated_genes | Top_51_to_100_upregulated_genes | Top_1_to_50_downregulated_genes | Top_51_to_100_downregulated_genes |
|---|---|---|---|
| Gm4070 | Slc25a4 | Serpina11 | Fgg |
| Sprr1a | Src | Gm4952 | Apoh |
| S100a11 | Myo5a | Cyp2c67 | Hgd |
| Myof | Peg3 | Cml2 | Cyp2c37 |
| Fbln2 | Abcc5 | 1100001G20Rik | Cps1 |
| Col1a1 | Cd24a | Orm1 | Mug2 |
| Serpine1 | Dbn1 | C8g | Serpinc1 |
| Ccl7 | Anln | Gck | Hal |
| Plat | Ahnak | Ces1b | Ces1d |
| Bicc1 | Emp1 | Gjb1 | Dpys |
| Ctgf | Slco3a1 | Uroc1 | Serpina3m |
| Map1b | Phlda3 | Elovl3 | Ces3a |
| Tes | Slc7a1 | Gm4788 | Angptl3 |
| Vim | Itpr3 | Cyp1a2 | Pon1 |
| Thbs1 | Ivl | Slc22a30 | Aox3 |
| Tnc | Spp1 | Gys2 | Cyp2f2 |
| Top2a | Ehd2 | Mup10 | Apoa2 |
| Xirp2 | Cldn4 | Cyp8b1 | Uox |
| Ltbp2 | Afp | Sdr9c7 | Cfhr2 |
| Mmp12 | Ampd3 | Serpina1d | Ces1c |
| Fgd3 | Cenpf | Cyp2d26 | F10 |
| Zfp462 | Fbn1 | Slc25a47 | Akr1c6 |
| Anxa3 | Abcc1 | Tdo2 | C8b |
| Arl4c | Nrk | Slc27a5 | Rgn |
| Krt19 | Col12a1 | Slc27a2 | Apoa5 |
| Ddr1 | Fads3 | Inmt | Slco1a1 |
| Jag1 | Afap1 | Pzp | Cyp2c29 |
| Clip2 | Ezr | Baat | C8a |
| Tgtp1 | Col5a2 | Proz | Cyp2d10 |
| Cpe | Ccl24 | Cyp2c44 | Pck1 |
| Phgdh | Ankrd1 | Pigr | Cyp3a11 |
| S100a6 | Tuba1a | Amdhd1 | Hrg |
| Basp1 | Abcc4 | Mup21 | Selenbp2 |
| Tagln | Ak1 | Cyp4a12a | Hpd |
| Sox4 | Serpinb9b | Ppp1r3c | Serpina3k |
| Mboat1 | Kif3c | Gnmt | G6pc |
| Cenpe | Ngfrap1 | Otc | Cyp2d9 |
| Loxl2 | Serpinb6a | Cpn2 | Rdh7 |
| Flna | Glipr2 | Cyp3a25 | Mup11 |
| A430105I19Rik | Shc2 | Hsd3b5 | 2810007J24Rik |
| Ccl2 | Usp18 | Tat | Plg |
| Gsta1 | Maff | Serpina3c | Ugt2b1 |
| Tgfb2 | Itga3 | Aldob | Apoa1 |
| Lgals1 | Ppp1r18 | Ces1e | Cyp2e1 |
| Flnc | Pdgfb | Cyp2a12 | Abcb11 |
| Prrg4 | Epdr1 | Upb1 | Mug1 |
| Epha4 | Plod2 | Haao | Slco1b2 |
| Slc25a24 | Lmcd1 | Ugt2a3 | Car3 |
| Cdh17 | Hells | Ces3b | Fabp1 |
| Cnn2 | Arhgap11a | Slc10a1 | Bhmt |

**Supplemental table 2. The 100 most upregulated genes and 100 most downregulated genes between 0-96h in RNA-seq_0h-192h dataset.**

### Supplemental Materials and method

Determining ideal K

To find the ideal number of K for each heatmap (Figures 1, 4, and 5), we calculated the within sum of squares and between sum of squares for K between 2 and 10. To find the within sum of squares, we calculated the centroid of the normalized EC/CE of each time point within each cluster in a k-means clustering. We then calculated the euclidean distance between each time point for each individual genes/areas of open chromatin and the centroid of the relevant time point. For each gene/area of open chromatin we calculated the euclidean norm and by squaring and summing the euclidean norm we calculated the within sum of squares for each cluster. The total within sum of squares for all clusters was calculated by summing the within sum of squares of all clusters. To find the between sum of squares, we first calculate total sum of squares of the normalized EC/CE. This is done in the same way as finding the within sum of squares for each cluster, but instead of using the genes/areas of open chromatin in each cluster, we use all of the genes/areas of open chromatin in the dataset. We then find the between sum of squares by subtracting the within sum of squares from the total sum of squares. A ratio was then calculated by dividing the between sum of squares by the within sum of squares and penalizing by dividing the ratio with the number of K clusters. A higher number designates a better fit to the data (Supplemental Figure 1).

RNA-sequencing

Cultured cells were singularized using trypsin (Thermo Fisher # 12605010). RNA was isolated using Qiagen RNeasy Mini Kit (Qiagen # 74104). Fetal liver sections were harvested and lysed before RNA was isolated. The RNA was sequenced using the following procedure. One hundred nanograms of total RNA was used to prepare sequencing libraries using the LM-Seq (Ligation Mediated Sequencing) protocol [24]. Final cDNA libraries were quantitated with the Qubit Fluorometer (Life Technologies, Carlsbad, CA) and multiplexed with 24–51 total indexed libraries per lane. The RNA-seq_0h-24h_EGFP dataset was sequenced using the HiSeq 3000 system (Illumina, San Diego, CA), and the RNA-seq_0h-192h and RNA-seq_0h-169h_EYFP was sequenced using the HiSeq 2500 system (Illumina, San Diego, CA). All datasets were sequenced using single end reads of 64 or 69 bp and an index read of 10 bp. The RNA-seq_Fetal_liver dataset was prepared using the MinAmp protocol as follows: We isolated fragmented rRNA depleted polyA+ RNAs using the Life Tech Ribominus Kit and 5X fragmentation buffer in addition to the Qiagen Oligotex Kit. This template was used to generate cDNA. Subsequent steps followed the Illumina Single Read preparation kit (Illumina, San Diego, CA). After the Illumina SR adapters were ligated, 200-300 bp DNA fragments were isolated via gel electrophoresis. Then fourteen cycles of polymerase chain reaction (PCR) were performed to amplify the selected fragments using the Illumina supplied PCR primers and protocol. The sample was quantitated with the Thermofisher (was Invitrogen) Qubit fluorometer (Q32857). The samples were sequenced using either the GAII or the GAIIx (Illumina, San Diego, CA).

ATAC-sequencing

ATAC-sequencing was performed according to the omni ATAC-seq protocol [25] with the following change: after isolated the nuclei were washed 3x in 100ul PBS. Then the

ATAC-seq samples were sequenced by the following procedure. ATAC-seq samples were normalized and multiplexed with one to four samples per lane. The Select-A-Size DNA Concentrator Kit (Zymo Research, Tustin, CA) double size selection protocol was followed as per the manufacturer's user guide to clean in addition to select for 100–700 base pair final libraries. Multiplexed lanes were quantitated with the Qubit Fluorometer (Life Technologies, Carlsbad, CA), and base pairs were confirmed with the Agilent Bioanalyzer High Sensitivity DNA Kit (Agilent, Santa Clara, CA). Lanes were sequenced as paired-end reads on the HiSeq 3000 system (Illumina, San Diego, CA).

Gene expression graphs

The graphs of normalized marker gene expression were created using the R packages ggplot2 (v.3.2.1) and gridExtra (v.2.3).

Histogram of distance between open chromatin and TSSs

The areas of open chromatin were annotated (annotate peak function from HOMER (V.4.11.1)) to either the TSSs of the nearest genes using mm10 or as a negative control, to an equal number of random locations on the genome. The data was binned based on the distances, and the histogram was plotted using ggplot2 with different bin sizes. Finally, we subtracted the histogram table of distances between the randomly selected genome areas and areas of open chromatin from the histogram table of distances between the TSSs and areas of open chromatin, allowing us to reduce the noise and identify the actual consensus pattern of openness around TSSs in our dataset.
